# Affordable and robust phenotyping framework to analyse root system architecture of soil-grown plants

**DOI:** 10.1101/573139

**Authors:** Thibaut Bontpart, Cristobal Concha, Valerio Giuffrida, Ingrid Robertson, Kassahun Admkie, Tulu Degefu, Nigusie Girma, Kassahun Tesfaye, Teklehaimanot Haileselassie, Asnake Fikre, Masresha Fetene, Sotirios A. Tsaftaris, Peter Doerner

**Affiliations:** Institute for Molecular Plant Science; School of Biological Sciences, University of Edinburgh, Edinburgh, UK; Institute of Digital Communications, School of Engineering, University of Edinburgh, Edinburgh, UK; Ethiopian Institute for Agricultural Research, Debre Zeit, Ethiopia; International Crops Research Institute for the semi-arid tropics, ICRISAT-Ethiopia, Addis Ababa, Ethiopia; College of Natural Sciences, Addis Ababa University, Addis Ababa, Ethiopia; Ethiopian Biotechnology Institute, Addis Ababa, Ethiopia; Ethiopian Academy of Sciences, Addis Ababa, Ethiopia

**Keywords:** Image-based plant phenotyping, root system architecture, soil-grown root systems, rhizobox, chickpea, Raspberry Pi, Phenotiki

## Abstract

The analysis of root system growth, root phenotyping, is important to inform efforts to enhance plant resource acquisition from soils. However, root phenotyping remains challenging due to soil opacity and requires systems that optimize root visibility and image acquisition. Previously reported systems require costly and bespoke materials not available in most countries, where breeders need tools to select varieties best adapted to local soils and field conditions. Here, we present an affordable soil-based growth container (rhizobox) and imaging system to phenotype root development in greenhouses or shelters. All components of the system are made from commodity components, locally available worldwide to facilitate the adoption of this affordable technology in low-income countries. The rhizobox is large enough (~6000 cm^2^ visible soil) to not restrict vertical root system growth for at least seven weeks after sowing, yet light enough (~21 kg) to be routinely moved manually. Support structures and an imaging station, with five cameras covering the whole soil surface, complement the rhizoboxes. Images are acquired *via* the Phenotiki sensor interface, collected, stitched and analysed. Root system architecture (RSA) parameters are quantified without intervention. RSA of a dicot (chickpea, *Cicer arietinum* L.) and a monocot (barley, *Hordeum vulgare* L.) species, which exhibit contrasting root systems, were analysed. The affordable system is relevant for efforts in Ethiopia and elsewhere to enhance yields and climate resilience of chickpea and other crops for improved food security.

**Significance Statement:** An affordable system to characterize root system architecture of soil-grown plants was developed. Using commodity components, this will enable local efforts world-wide to breed for enhanced root systems.

## Introduction

The distribution of plant roots is referred to as root system architecture (RSA), which changes over time as the plant grows and adapts to soil conditions (de Dorlodot *et al*., 2007, Tian and Doerner 2013). RSA describes and quantifies parameters such as the shape, extent and density of the root system, and hence, is a major descriptor of plant resource acquisition capacity and ensuing competitive success. Often, RSA analysis is reduced to parameters of *topology*, which is a generalized representation of individual root hierarchy resulting from growth; distinguishing the main root (primary root, PR), which develops first, from which branches (secondary root, SR) emerge, with further branches emerging from the SR (tertiary root, TR) and so on (Lynch 1995). Another set of traits frequently analysed are related to individual root *morphology*, for example root diameter. (Lynch 1995). Most dicotyledonous plants develop a root system in which roots can easily be parsed in this manner, whereas monocotyledonous plants exhibit a more complex system that defies simple hierarchical characterization as they lack a clear main root (Smith and De Smet 2012, Atkinson *et al*., 2014).

Roots not only provide anchorage but also acquire resources (water and nutrients) and for both of these functions, the *spatial distribution* of the root system within the soil is a critical determinant of successful exploitation of below-ground resources, most of which are non-uniformly distributed. Therefore, topology and morphology are necessary, but insufficient descriptors of root systems. Root system growth in the soil gives rise to *emergent* system parameters, including, but not limited to the convex hull, the centroid, local root density and the total length of the root system. The convex hull is defined as the area of the smallest polygon, with interior angles less than or equal to 180°, covering the whole root system, when projected onto a 2D plane (Pound *et al.*, 2013), while the centroid identifies the centre of mass of that geometric shape. Root density, which relates to the intensity of soil exploration, is the length or area of roots in a given area or volume; and the total length or area of the root system is a good proxy for the root biomass produced by the plant.

Such RSA descriptors can be used for comparative purposes (Kutschera and Lichtenegger 1997, Bouma *et al.*, 2001, Pages 2016) and, because they relate to root system function, can inform crop improvement (Burridge *et al.*, 2016, Burridge *et al.*, 2017, Morris *et al.*, 2017). Emergent RSA parameters relate to RSA at any moment in time, and extending this analysis to a time series also reveals important features such as global or local growth and branching rates that inform on the plant’s resource capture and internal resource distribution strategies. The RSA of a plant is the result of a genetically determined general pattern that is modified by the specific soil and rhizosphere conditions that it experiences during its life. Thus, RSA is plastic and adaptive, and its analysis is relevant for the improvement of plant performance, specifically in extreme conditions. For RSA analysis to contribute to plant improvement efforts, it is desirable to be simple, high-throughput and placed into the hands of those that require this information to develop elite lines (Lynch 1995, Lynch 2007, Burridge *et al.*, 2016).

Soil opacity is a major problem when studying plant RSA. Modern techniques such as X-ray computed tomography (Heeraman *et al.*, 1997, Morris *et al*., 2017) allow the three-dimensional (3D) reconstitution of the root system, even when grown in soil, but are slow, have limits on the soil volume that can be sampled and require prohibitively expensive equipment. For this reason, many lab-based growth systems have been developed that are geared toward visualising root systems and their RSA by visible wavelength imaging, including: growth matrices such as gellan gum (Iyer-Pascuzzi *et al.*, 2010), transparent synthetic soil (Downie *et al.*, 2012) and hydroponics (Mathieu *et al.*,2015). In these cases, roots are not developing, growing or interacting with the natural abiotic and biotic environment in which they evolved (Morris *et al.*, 2018), and there is strong evidence that root growth behaviour in these systems is different from soil-grown plants (Rellan-Alvarez *et al*., 2015, Silva-Navas *et al.*, 2015). Hence, when developing a plant growth system to study RSA parameters, one faces a trade-off between realistic growth conditions and root visibility.

Many investigators have tried to combine ease of root system detection and growth in the soil substrate. The first investigator known to have developed a “root box” was Julius von Sachs in the 19^th^ century (Sachs 1865, Kutschera 2015). Since then, many soil-based growth systems, referred to as rhizoboxes, have been developed, several of which allow observation of only a small fraction of the root system growing in 3D by introducing transparent tubes into the soil (Sanders and Brown 1978). Other soil-based growth systems provide relatively thin layers of soil bordered by one or two transparent surfaces to visualise roots pressed against them, thus collapsing a variable fraction of the entire root system in 3D against a transparent surface for a 2D representation (Neumann *et al.*, 2009). Such 2D systems have been reported for the dicotyledonous species *Arabidopsis thaliana* (Devienne-Barret *et al.*, 2006, Rellan-Alvarez *et al.*, 2015), tomato (Dresbøll *et al.*, 2013, Rellan-Alvarez *et al.*, 2015), lupine (Leitner *et al*., 2014), sugar beet (Bodner *et al.*, 2017), or monocots such as rice (Price *et al.*, 2002, Shrestha *et al.*, 2014), and wheat (Jin *et al.*, 2015). These systems have allowed testing of plant growth behaviour in waterlogging (Dresbøll *et al.*, 2013), low moisture stress (Avramova *et al.*, 2016, Durand *et al.*, 2016) or contrasting nutrient availability conditions (Jin *et al.*, 2015). With the exception of two previous studies (Shrestha *et al.*, 2014, Jin *et al.*, 2015), the growth systems for such studies were ≤ 1m in height (Rellan-Alvarez *et al.*, 2015, Avramova *et al.*, 2016), which limited analyses to early growth phases of most plants. Although the constrained growth in such containers is distinct from plant growth in the field, the results obtained are informative and robotic platforms for 2D growing root phenotyping have been developed (Nagel *et al.*, 2012, Wu *et al.*, 2018). A common drawback of these systems is their substantial cost caused by use of bespoke or expensive components which precludes their use at larger scales or implementation in low-income countries.

2D growth systems enable the acquisition of root system images using flat-bed scanners (Devienne-Barret *et al.*, 2006), CCD camera(s) (Rellan-Alvarez *et al.*, 2015) or neutron radiography (Leitner *et al.*, 2014). To quantify and analyse RSA parameters numerous software packages such as Smart-Root (Lobet *et al.*, 2011), GLO-RIA (Rellan-Alvarez *et al.*, 2015) or Root System Analyzer (Leitner *et al.*, 2014), have been developed. However, many are based on destructive analysis, or require artificial substrates and require significant human intervention by highly-trained operators (Kuijken *et al.*, 2015).

In this study, we describe a large-dimensioned (150 x 45 cm), simple and affordable rhizobox developed to grow plants in soil and analyse their changing patterns of RSA. The system comprises commodity components that are readily sourced in most parts of the world. Our rhizobox was optimised to observe a large fraction of the root system in 2D. We designed and built an imaging station, also based on affordable commodity components, permits the acquisition of high-resolution (approx. 9000×2700 pixels) images using low-cost cameras. RSA parameters such as root system growth rate, shape, extent and density were quantified. To evaluate its robustness and utility in different environments, the rhizobox system developed in Edinburgh, United Kingdom was also established and tested at the Debre Zeit Agricultural Research Centre, Bishoftu, Ethiopia using two chickpea cultivars commonly grown in Ethiopia.

## Results

### Establishment of a commodity component-based system to visualise soil grown plant root systems

A modular, commodity component-based rhizobox was designed, assembled (Figure 1A, B) and evaluated for chickpea and barley growth in soil (Figure 1C). One plant was grown per rhizobox in a 6 mm-thick layer in a soil volume of ~3.7 dm^3^ on supporting racks (Figure 1C). Growth and root system distribution were imaged every 2-5 days until the bud-filling stage for chickpea with an imaging station that contained five Raspberry Pi cameras (Figure 2). Raw image data was processed in a pipeline to assemble composite images for each root system at a given time point (Figure 3). The following emergent root system parameters were analysed: Total Area, Convex hull area, Total Length, Growth rate, Depth, Width, Centroid, Solidity and Density (Table 1). Detailed information for rhizobox components and assembly, use and data capture are presented in Material and Methods and in Supporting Information.

**Figure 1.**
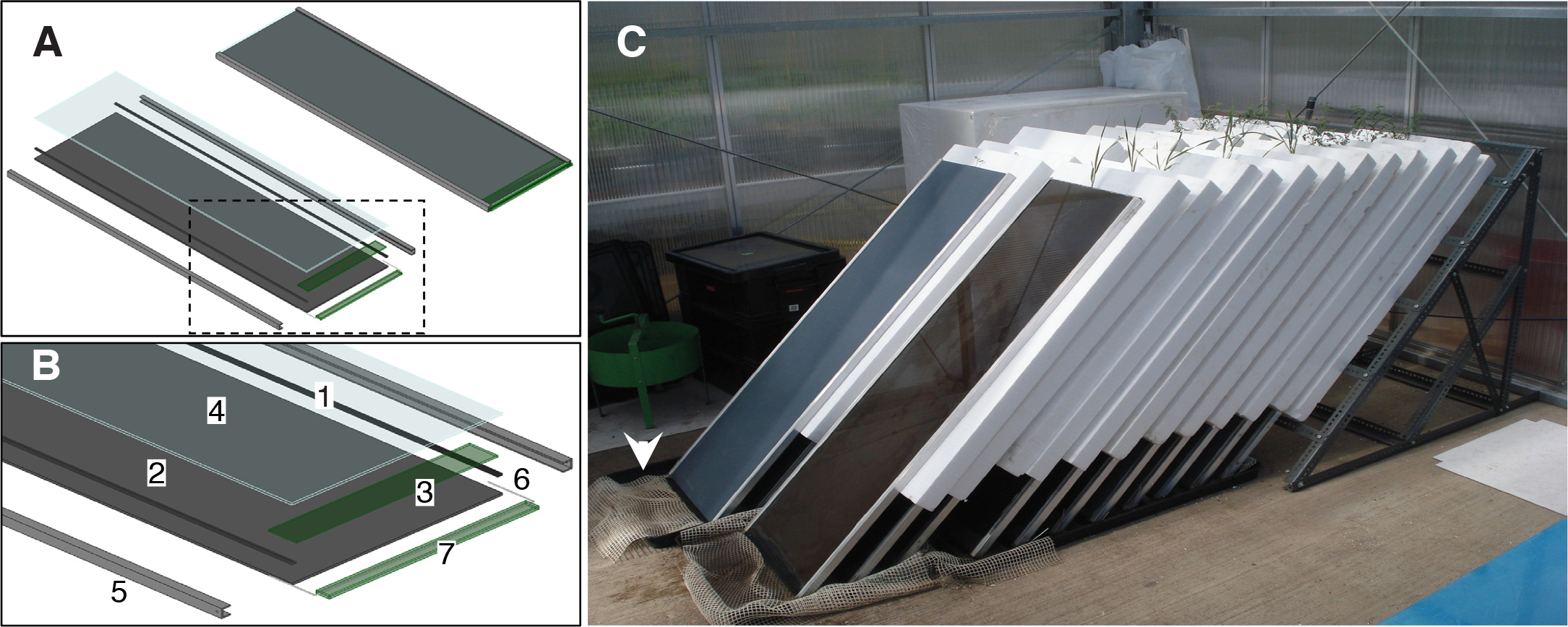
Rhizobox components and support. Exploded **(A**, left) and closed **(A**, right) view diagram of a rhizobox designed with FreeCAD 0.16. The enlarged exploded view **(B)** shows silicon strips (1) glued along the PVC sheet (2) laid horizontally. An inner piece of nylon mesh (3) is inserted at the base before soil loading. The glass pane (4) is added after soil loading. Two aluminium U-channels (5) linked by a steel wire (6) inserted in the outer nylon mesh (7), make up the frame that closes the rhizobox. (C) Rhizobox support. 22 rhizoboxes are aligned in two rows and separated with white polystyrene sheets. Rhizobox glass side (anterior) faces downwards with an inclination of 45°. Pieces of anti-slip mesh were laid in trays holding the rhizoboxes (arrowhead).

**Figure 2.**
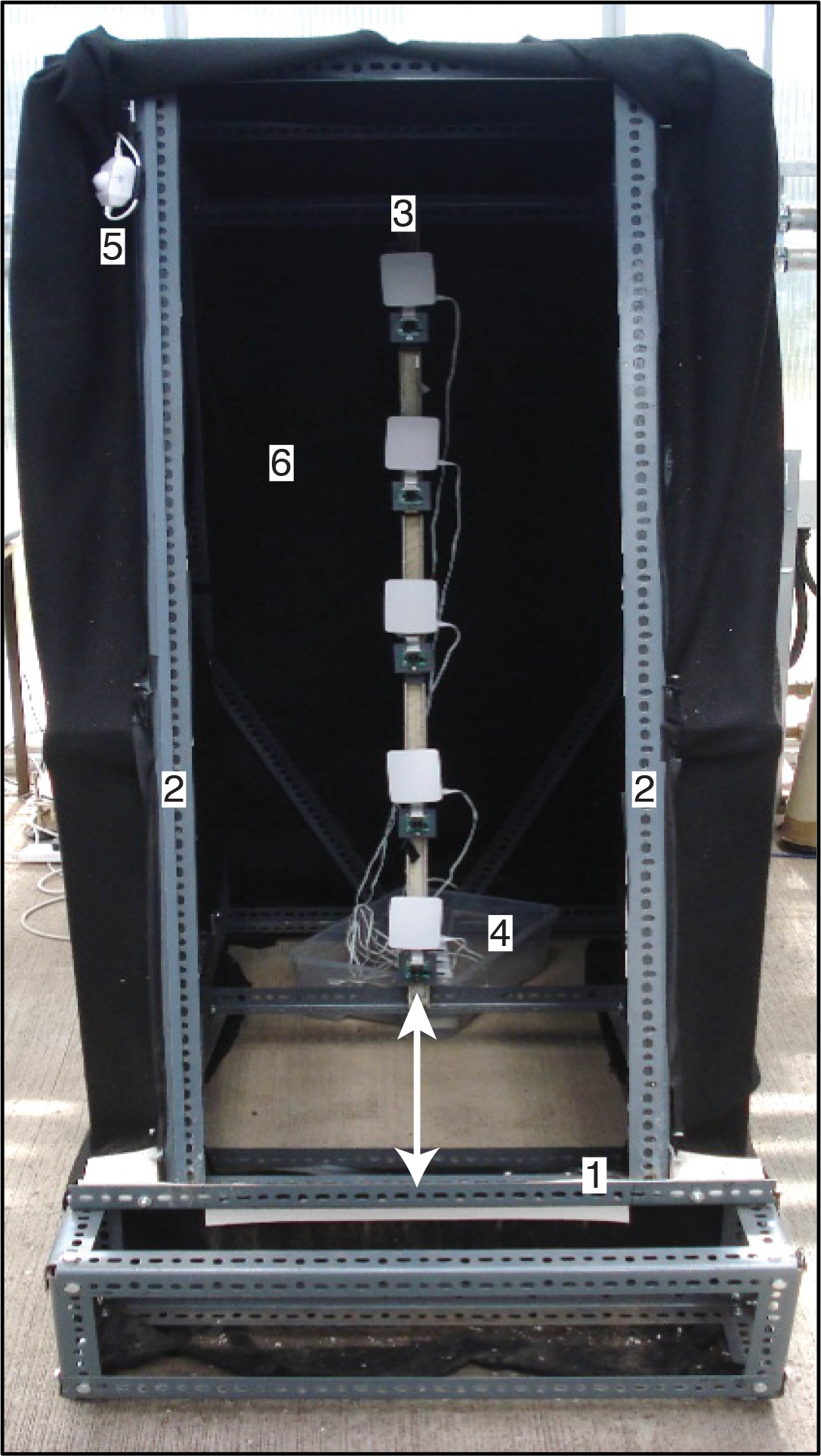
Imaging station for imaging of a rhizobox. The imaging station is constructed from slotted angle and holds a rhizobox, placed on a step (1) padded with a silicon sheet, at a 15° inclination, when placed against two lateral bars (2). An internal camera array (3), is placed parallel to the rhizobox, at a distance of 78 cm (indicated by the double-headed arrows). All cameras are plugged to a common multi-socket (4) and an external light switch (5) controls LED-lighting intensity inside the imaging station. The whole structure is covered with black felt (6) to suppress stray external light.

**Figure 3.**
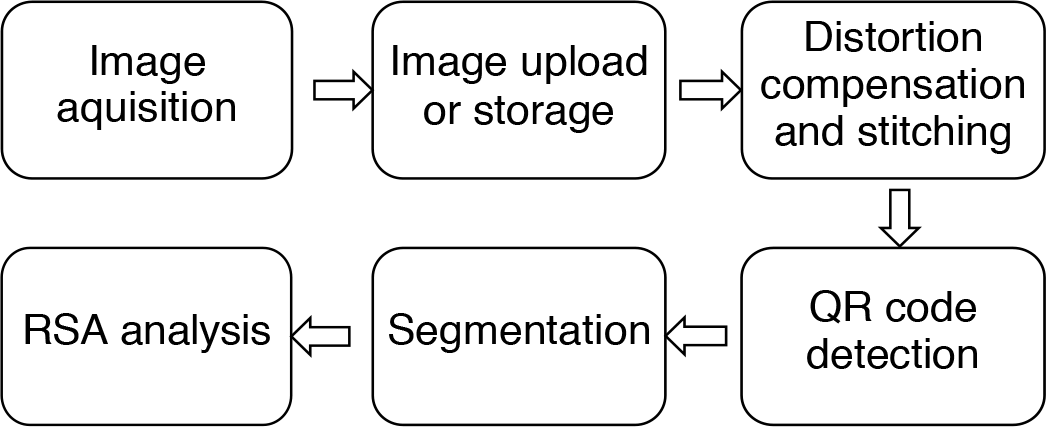
Diagram of image capture and processing pipeline. Images are acquired *via* Phenotiki interface, which can be automatically uploaded to the cloud. The interface also allows the user to download all the acquired images, in case of suboptimal Internet connection. For each rhizobox, five images are corrected for distortion, compensated and stitched together to form a single, un-distorted image of the entire anterior face of the rhizobox. QR code detection provides unique identification of each assembled rhizobox image. The segmentation step extracts the root system from soil background. Root system architecture (RSA) is analysed from the segmented root system with customized algorithms.

**Table 1.** Root system architecture parameters. All parameters were converted from pixels to centimetres.

| Parameter | Unit | Computing method |
| --- | --- | --- |
| Total Area | cm<sup>2</sup> | Number of pixels detected as root |
| Convex hull area | cm<sup>2</sup> | Area of the smallest convex hull enclosing the root system |
| Total Length | cm | Number of the pixels of the skeletonized root system |
| Growth rate | cm<sup>2</sup>.d<sup>-1</sup> | Difference of root system area over unit of time |
| Depth | cm | Maximal vertical extension |
| Width | cm | Maximal horizontal extension |
| Centroid | cm | Coordinates of the centre of mass with respect to the root system |
| Solidity | N/A | Total root area relative to convex hull area |
| Density | N/A | Number of root pixels within a defined square region |

### Optimal parameters to maximise the visual fraction of the root system

To optimize the visible fraction of the root system on the (front) glass side and minimize data loss caused by roots not growing against this side, we built modified rhizoboxes by replacing the PVC sheet (back) with a second glass pane (termed double glass rhizoboxes), for which we could capture images from both sides to reliably assess the fraction of the root system exposed to the back side.

We tested two soil compression methods: against the side facing down (anterior compression, against front glass side) or against the side facing up (posterior compression, against back side; normally PVC, here glass side), as well as two inclination angles, 30° and 45° from the vertical (Figure 4A). At 48 days after sowing (das), we took images of both sides, which were then segmented to delineate root from soil. The overlap of front and back sides (Figure 4B, 4C) illustrates the visibility of root system in the worst and best cases observed for anterior 30° and posterior 45° conditions, respectively. The data were analysed (Figure 4D-4F): the percentage of root pixels counted on each side (front in blue, back in orange) compared to total pixels counted per rhizobox (n=4 for each condition, Figure 4D); mean percentage of the four rhizoboxes per condition (Figure 4E). With compression against the anterior side and 30° inclination, only 42% of the roots were observed on the front side. At 45° inclination, 53.5% of the root system was visible on the front side. When the soil was compressed against the posterior side, the fraction of the root system visible on the front side reached a mean of 73.3% and 75.4% at 30° and 45° inclination, respectively. The lowest front side root system visibility of the rhizobox at 45° inclination was 69.7% whereas it was 59.3% for the rhizobox at 30° inclination (Figure 4F). The posterior 45° condition resulted in higher maximal front visibility (86.2%), compared to a maximum of 79.2% for the posterior 30° condition.

**Figure 4.**
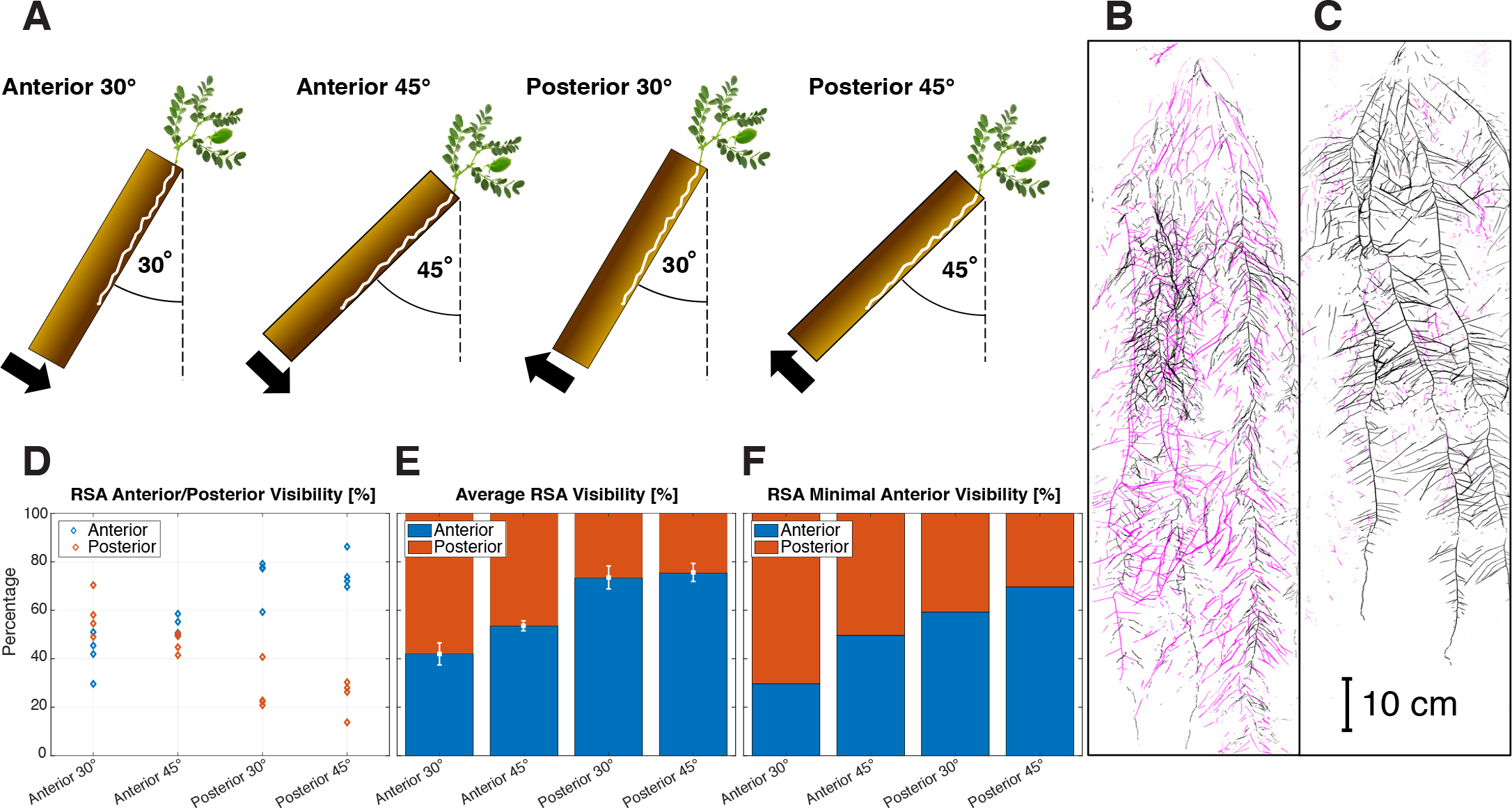
Evaluation of root system growth with double glass rhizoboxes. **(A)** Diagram of rhizoboxes with soil compressed against the anterior or posterior side, inclined at 30 or 45° from vertical, respectively. The black arrow indicates the direction of soil compression. Brown rectangles represent soil in rhizoboxes. Images were acquired 48 days after sowing. Each side of a double glass rhizobox was imaged and overlapped to compare the root part visible on the anterior (in black) and posterior side (in magenta). **(B)** The worst case in condition Anterior 30°. **(C)** The best case in condition Posterior 45 °. **(D)** Percentage of roots visible on anterior (blue) and posterior (orange) sides for each rhizobox in the four conditions. **(E)** Average % of the root visible on each side ± standard error (n=4). **(F)** The minimal % of root visible for each condition. For **(D-F)**, the number of pixels corresponding to roots visible on each side was determined after segmentation.

Although these differences (between posterior compression at 30° and 45°, respectively) in the observable (front) fraction of the root system were not statistically significant (p=0.74), we concluded that compression against the posterior side and 45° inclination was the best condition to maximize the visible fraction of the root system in the rhizobox. Therefore, all other rhizoboxes in this study were prepared in this manner.

### Root system analysis

Images of four chickpea plants grown in rhizoboxes in Edinburgh were acquired three times per week from 10 until 35 das with 2–5-day intervals (Figure 5A, Movie S1). This time course included the onset of flowering in these chickpea plants (31-33 das); vertical growth was not restricted as the roots did not reach the bottom of the rhizobox. The segmented root system at 10, 17, 24, 31 and 35 das for one of the four rhizoboxes clearly shows that root system growth persists beyond the onset of flowering at 31 das (Figure 5A). An overlap of the convex hull for each time point mapped on the final segmented root system at 35 das reveals that most growth activity at this late time point occurs in the lower half of the root system to increase its lateral extent (right hand panel, Figure 5A).

**Figure 5.**
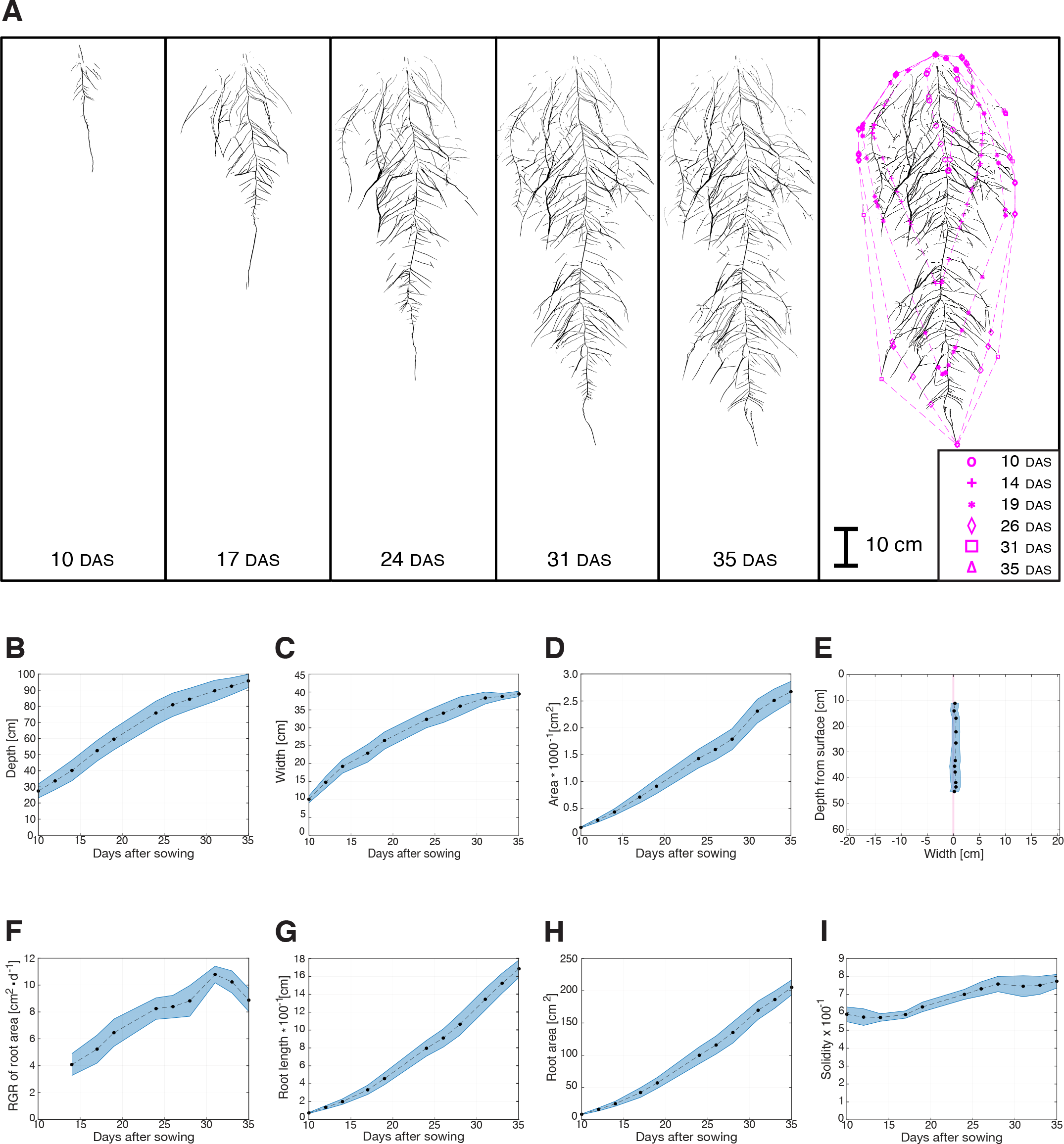
Evolution of root system parameters during chickpea development in rhizoboxes. **(A)** Segmented images of the root system in one rhizobox at 10, 17, 24, 31 and 35 days after sowing (das). The convex hull corresponding to each time was overlapped on the segmented root system image at 35 das. **(B-I)** Root system parameters analysed from four rhizoboxes. The dashed line shows mean and the shaded region indicates standard error. **(B)** Depth: maximal vertical extension. **(C)** Width: horizontal distance between the leftmost and rightmost root system pixel. **(D)** Convex hull area: the smallest polygon, with no interior angles less than 180°, covering the whole root system. (E) Centroid: coordinates of the centre of mass with respect to the root system. Each data point represents the root system’s centroid (from top to bottom) at 10, 12, 14, 17, 19, 24, 26, 28, 31, 33 and 35 das, respectively. **(F)** Relative growth rate: difference of root system area at time *t*_*n*_ relative to time *t*_*n-2*_ over time. **(G)** Total length: quantified as the number of pixels of the skeletonised segmented image. **(H)** Total area: computed as summed root pixels. **(I)** Solidity: calculated as the ratio of root area over convex hull area.

Segmented images were analysed to extract the emergent RSA parameters and plotted over time (Table 1, Figure 5B-I).

Root system depth (Figure 5B), which corresponds to the vertical position of the deepest root pixel, increases linearly during that time course, with an inflection observed at 26 das, reflecting reduced root apical growth. The maximal value recorded for that parameter was ~1 m, confirming that in our system root growth was not restricted in terms of depth. Root system width (Figure 5C), computed as the horizontal distance between the right- and left-most pixels, also increases, but is likely to be constrained by the rhizobox dimensions from 31 das onwards.

Convex hull area (Figure 5D) increases linearly along the time course to reach ~2750 cm^2^ after 35 days. The average location of all the root pixels (centroid) was plotted relative to the coordinates of the rhizobox. On average, the centroid is aligned to the central vertical axis, showing that the root system is equally spread along the horizontal axis (Figure 5E). Smoothed root system area growth rate (Figure 5F) was computed as the ratio of the difference between the root system area at time *t*_*n*_ and at time *t*_*n-2*_, over the difference in time (Δdas). This growth rate initially increased over time then decreased around flowering time (31 das). Total length of the root system (Figure 5G) was calculated as the number of pixels after segmented images were skeletonized (Giuffrida *et al.*, 2015, Wu *et al.*, 2018); this parameter increased exponentially over time during the experiment, reflecting the increased number of actively growing root tips. The total area of the root system (Figure 5H), computed by counting the number of pixels segmented as *“*root*”* in an image, exhibits a similar curve profile as root system length over time, these two parameters are well correlated (R^2^=0.99, Figure S1). Root system solidity (Figure 5I), the ratio of total root area over the convex hull area, decreased slightly until 14 das, then increased between 17 and 28 das to finally reach a saturation phase. Root system solidity is positively correlated with total root area, albeit not as strongly (R^2^=0.45, Figure S2), suggesting changing growth and resource capture strategies over time.

We analysed dynamic changes to root distribution within the convex hull by assessing fractional root area in a non-overlapping square grid of ~3 cm, visualised as heatmap of root densities within the associated convex hull (Figure 6). The distribution of root pixels per unit area within the convex hull reveals that chickpea roots fill the root system area (convex hull) relatively evenly (Figure 6A), with no clear emphasis in any one area of the system. This is also apparent when examining differential growth where the difference in root pixels for each time *t*_*n*_ and time *t*_*n-1*_ is plotted (Figure 6B). This analysis reveals a combination of two growth modes: *i)* expansion of total area (convex hull) sampled to contact new soil volumes and *ii)* growth within the convex hull to increase the plant-soil interface (Figure 6B). Moreover, around the time of flowering and after (24–35 das), growth in deeper soil strata is emphasized, and reduced in strata close to the surface.

**Figure 6.**
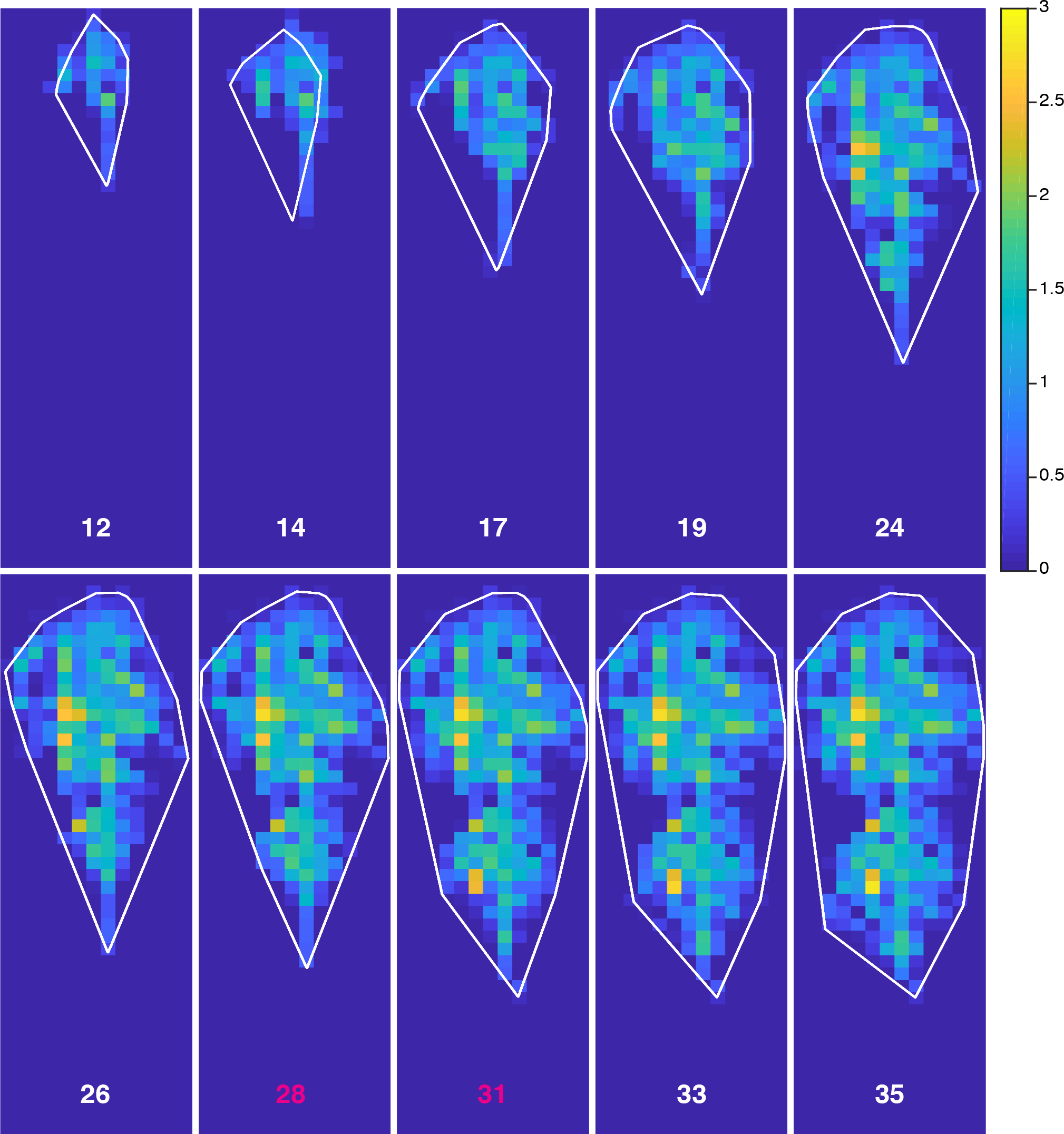
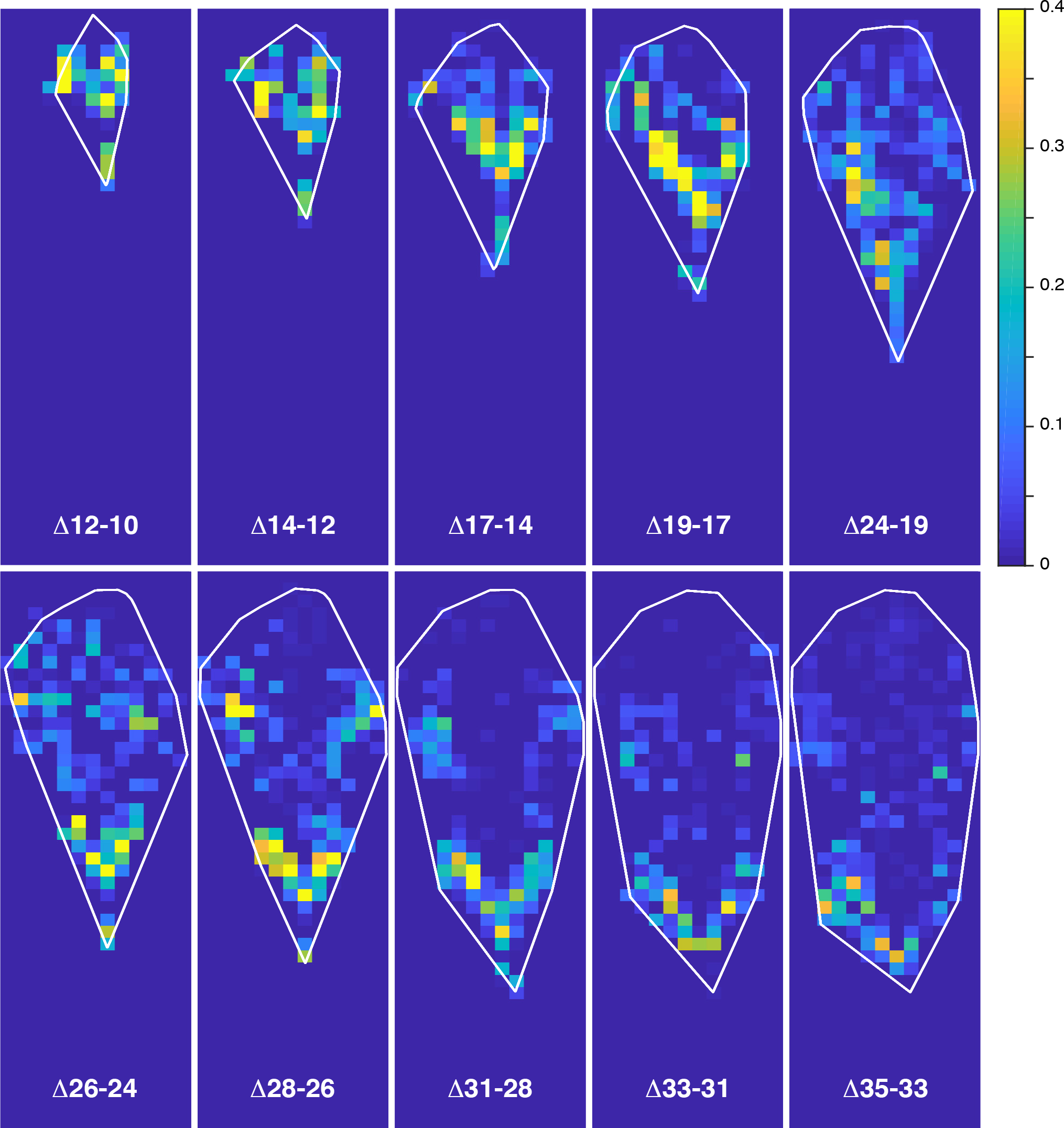
Chickpea root area density. That parameter was computed in a grid of 200 x 200 pixel squares covering the whole rhizobox. The root system corresponds to the chickpea shown in Figure 4, and days after sowing (das) are indicated. The scale unit corresponds to the number of root pixels per square. (A) Absolute root area density. (B) The rate of root area density change was computed as the difference of root pixels between time *t*_*n*_ and time *t*_*n-1*_, over the difference in time (Δ das).

### Comparability across sites and species

To evaluate whether the rhizobox system could be used for other, particularly monocot crop species, we tested the growth of barley. Barley (*var*. Concerto) was grown in rhizoboxes and the root system was imaged repeatedly until 25 das to test our approaches for segmentation and extraction of RSA parameters. These images were successfully segmented, which enabled all RSA parameters to be quantified over time (Figure S3). The detection of barley main roots was as robust as for chickpea, although some thinner lateral roots are not always detected. Monocot root systems often exhibit thinner secondary or tertiary roots that challenge the segmentation process established for chickpea. Most RSA parameters (width, length, area, convex hull area) increased exponentially in barley whereas depth and root growth rate increases were linear (Figure S3). In contrast to chickpea, root solidity in barley decreased continuously indicating that convex hull area increases faster than root area. When analysing local root densities, growth activity of the root system until 25 das is focussed on accessing deeper soil strata (Figure S4A,B).

The rhizobox developed in Edinburgh (UK) was also tested in Debre Zeit (Ethiopia). All commodity components for rhizobox construction at Debre Zeit were sourced in Ethiopia, except for silicone spacers. As the majority of chickpea field cultivation in Ethiopia is conducted in vertisols, a unique, clay-rich soil, we aimed to test whether rhizoboxes would support chickpea growth and allow root system visualisation when filled with vertisol. When loaded with vertisol from local farmland, the rhizoboxes supported chickpea growth (Figure S5). Two local cultivars were tested: ‘Dhera’ (Kabuli variety, ICARDA FLIP 0163) and ‘Teketay’ (Desi variety, ICRISAT ICCV-00104). The main part of the chickpea root system is clearly visible as the local vertisol provides good contrast for segmentation (Figure S5).

## Discussion

We developed a growth (rhizobox) and imaging system using low-cost commodity components to study root development in soil non-destructively and at multiple times during the plant life cycle. The system is based on frugal engineering principles and can be operated without extensive training; therefore, it can be assembled and operated in most low- and middle-income countries. Images are acquired *via* the extended Phenotiki platform, stored and analysed. We routinely used this system to capture RSA parameters for crops, *e.g.* chickpea, barley. Despite its affordability, we can here document root growth over an area of 6000 cm^2^ (150 x 40 cm) in images of approximately 9000×2700 pixels, providing *ca.* 24 megapixels. Low-cost commodity components and the features of the data acquisition and processing pipeline will now enable this powerful tool to be used by breeders in many countries to inform their strategies at enhancing crop performance.

### Growth system validation

To reflect genetically encoded (genotype) and adaptive root growth behaviours (environment) with higher fidelity, and generate tools informative for breeders, we would focus on soil-grown roots. For example, a recent study characterising root systems of 270 chickpea genotypes in a semi-hydroponic system (Chen *et al*., 2017), showed that varieties exhibiting short root systems in soil (e.g. ICC 283 and ICC 1882; Kashiwagi *et al*., 2005) display intermediate or deep root systems in semi-hydroponic growth. Hence, non-soil based systems lack predictive value for breeders: in those conditions, roots do not face soil physical constraints, experience a completely different hydrology and nutrient distribution, and do not interact with the microbiome, all factors to which root systems are known to respond adaptively. We conclude that soil-based growth systems reflect root behaviour in natural conditions with higher fidelity, and therefore let breeders focus their efforts on the most promising germplasm.

Soil-filled systems with one transparent side for root system visualization have previously been developed (e.g. Sachs 1865, Price *et al*., 2002, Devienne-Barret *et al*., 2006, Bodner *et al*., 2017). These systems differ in their dimensions, and hence, the volume of soil available for root system growth. Small growth systems with extremely thin soil layers that can be readily handled have been reported, but these are too small to support unimpeded root growth throughout the life cycle of most crops (Rellan-Alvarez *et al.*, 2015). However, thicker soil layers or double glass large-dimensioned rhizoboxes make regular handling challenging while costs increased considerably, thereby the ability to process large numbers of rhizoboxes is reduced and thus overall throughput hampered. The system reported here balances large overall dimensions, which impose few constraints on root architectural development, with ease-of-handling requirements and low costs.

For maximum visualisation of roots, their gravitropism is usually exploited by inclining the growth box at angles between 15 to 45° from the vertical. We aimed to maximise the part of the root system visible on the imaged front glass side. Double glass rhizoboxes were used to assess how inclination angle and direction of soil compression affect the root system fraction visible on the front side. Even if the soil layer in the rhizoboxes described here is relatively thin (6 mm) compared to other systems (e.g. Nagel *et al.*, 2012, Jin *et al.*, 2015); a variable fraction of the root system will be hidden within it, or growing against the back side. The latter will only be seen in double glass rhizoboxes. Soil compression against the posterior side combined with an inclination of 45° was the optimal approach to maximise root system visibility on the anterior side (Figure 4A.). On average of 75.4% and a maximum of 86.2% of the visible root system was imaged on the anterior side (Figure 4D-F). When compared to a previous study (Nagel *et al.*, 2012), where the length of the root system imaged on the transparent side was compared to the length of the entire root system after soil removal, our system performs comparably to their best results: Nagel and colleagues (2012) report species-dependant differences in anterior visibility, ranging from 17% for *Zea mays* to 77% for *Arabidopsis thaliana.*

Moreover, a comparison of our system (6 mm soil thickness) with one containing more soil (34 mm soil) shows that a phylogenetically close relative to chickpea, faba bean *(Vicia faba*L.), does not perform markedly better with ~6 times more soil to exploit (Belachew *et al.*, 2018): on average at 28 das, root system depths were 72 and 84 cm, and convex hull areas were 2154 and 1789 cm^2^, for faba bean (Belachew *et al.*, 2018) and chickpea (this work), respectively. We conclude that our affordable growth and imaging system yields results very similar to those obtained with substantially more expensive and complex systems, *e.g.* automatic root phenotyping platform (Nagel *et al.*, 2012, Belachew *et al.*, 2018).

### Analysis of root system topology or root system architecture?

The extent, density, and rate of development of root systems reflect a plant’s ability to capture resources from their soil environment (Lynch 1995, Lynch 2007, Tian and Doerner 2013). Topological analysis of root systems, *i.e.* measuring PR length, the number, length and connectivity of SR and TR, does not comprehensively describe root system interactions with the environment and resources acquisition. By contrast, root system architecture (RSA), defined as the spatial distribution of plant roots in the soil, is an emergent property that includes additional parameters to describe the spatial interaction of roots with their environment. Lynch (1995) clearly distinguished root architecture from morphology *(e.g.* root diameter) or topology *(e.g.* number and hierarchy of SR), arguing that the latter two are not sufficient to describe root architecture, whereas the former can be used to derive the other two.

We developed our system primarily to empower local breeders to enhance crop performance by improving RSA. We reasoned that capturing architectural parameters such as depth, width, density, local growth rate, etc. which relate to the root system’s functional exploitation of soil resources, would provide actionable information for breeders, with the added benefit of being computationally less intensive than topological parameters.

### Robustness of analysis and significance of parameters

In any soil-based system that is non-destructively and multiply analysed by visible wavelength imaging, topological parameters are inherently incompletely captured. This is due to gaps in the imaged roots caused by soil obscuring parts of or entire roots. By contrast, architectural analysis is more tolerant to uncertainty and noise, as not every root or part of root must be identified for meaningful information to be obtained. Topological analysis would have been limited by hidden parts of the root system, resulting in discontinuous root topology. Nonetheless, a promising recent approach has demonstrated that such hidden root parts can be recovered with the use of deep neural networks (Chen *et al.*, 2018).

The focus on architectural parameters in the analysis of rhizobox images allowed us to develop algorithms to automatically extract parameters (*e.g*. convex hull area, root density) with direct utility for breeders and improve the efficacy of selection. RSA parameters reflect soil resource acquisition strategies: root system length and area describe the individual investment in root biomass for resource foraging and acquisition. During the vegetative phase, the continuous increase of root system length we observed for chickpea (Figure 5G) is commonly reported for other plants species grown in 2D soil-filled system (Price *et al.*, 2002, Devienne-Barret *et al.*, 2006, Leitner *et al.*, 2014, Yuan *et al.*, 2016). Root solidity (ratio root/convex hull areas) reflects a trade-off between foraging and space occupancy. It provides insight into the strategy to acquire soil resources: extend or intensify, with higher solidity reflecting more intense resource foraging within the explored area. Solidity in the chickpea cultivar tested shows a slight decreasing trend until 14 das (Figure 5I), implying that convex hull area increases faster than root system area. Resource foraging then becomes more intense between 14 and 28 das, as solidity increases. After 28 das, solidity is stationary as both convex hull area and root system area increase at a similar rate. These dynamics could indicate that the plant reduces its investment to intensive foraging, possibly because immobile resources within the densely rooted volume have been depleted, or its resource requirements and metabolic budgets have shifted priorities with the onset of reproduction (Koelewijn 2004).

Interestingly, the initial study of barley RSA revealed a different resource acquisition strategy to chickpea. The barley root system grows more extensively (predominantly in depth) rather than intensively, leading to a continuous decrease in solidity over the sample time. Consistent with this, highest root growth activity is consistently close to the deepest root tips (Figure S4B). Our rhizobox system allows inter-species comparisons which are relevant ecologically and for the development of novel multi-species or multi-cultivar cropping systems aimed at minimising competition for resource acquisition within a given environment (Li *et al.*, 2014, Wang *et al.*, 2014, Weiner *et al.*, 2017).

Studies in our group are in progress to utilize these tools to examine different germplasm for variations to these dynamics, their responses to limiting mobile (*e.g*. water and nitrogen), immobile (e.g. phosphate and iron) resources, interactions with the soil microbiota and associated changes to metabolism and physiology.

### Informing chickpea breeding in low- and middle-income countries

Previous studies with chickpea focussed on root topological parameters and were destructive (Krishnamurthy *et al.*, 1998, Serraj *et al.*, 2004, Ali *et al.*, 2005, Kashiwagi *et al.*, 2005, Kashiwagi *et al*., 2006, Pang *et al*., 2011, Purushothaman *et al*., 2017, Pang *et al*., 2018). In contrast, the rhizobox system reported here permits repeated segmentation and thus permits continuous observation of root system growth dynamics in a non-destructive way. The latter is a key feature for this system’s utility: for example, drought conditions are thought to be particularly damaging around the time of onset of flowering. Therefore the ability to analyse root system architectural parameters over time is crucial to identify germplasm that directs rapid, early and deep root growth to access residual water (Gaur et *al.*, 2008, Upadhyaya *et al.*, 2012).

This rhizobox system is currently being established and tested at the Ethiopian Institute for Agricultural Research (EIAR) at Debre Zeit (Bishoftu) in Ethiopia. Initial results (Figure S5) indicate that the system can be utilized with local vertisol soils, which are among the most challenging soils for agriculture due to their rheological properties (Jones *et al.*, 2013).

We conclude that the newly developed rhizobox system based on commodity components and powerful analytical tools will be useful to inform local breeders to address food security challenges by accelerating the enhancement of RSA-based traits associated with increased resilience and resource acquisition.

## Materials and Methods

### Rhizobox design and construction

The rhizobox (Figure 1A, B), made of the components described in Table S1, holds a 6 mm layer of soil between a sheet of polyvinylchloride (PVC; 1500 mm x 450 mm x 6 mm), a 6 mm silicone spacer, and a glass pane of the same dimensions as the PVC backing for a total soil volume of ~3.7 dm^3^. The assembly is held together by two aluminium U-channels on the sides, and a wire inserted into a folded piece of nylon mesh to close the bottom of the rhizobox. See Supplemental Methods for more details.

John Innes No.1 Young Plant Compost (Westland, UK) is sieved, and set to 50% (w:w) water content, then loaded horizontally into the rhizobox. The soil is manually spread uniformly, then compressed to ensure that the surface is level with the silicon strips. After adding the glass pane, the system is closed with the U-channel frame described above. See Supplemental Methods for more details.

### Plant material and growth conditions

In Edinburgh, chickpea *(Cicer arietinum* L.) seeds were imbibed by soaking in water for 18 h, then sown in John Innes No.1 compost (50 % (w:w)) for 2-3 days before transplanting the seedling into a rhizobox. In Ethiopia, chickpea seed were imbibed and sown directly into rhizoboxes filled with local vertisol. Barley *(Hordeum vulgare* L., variety *Concerto*, SRUC, UK) seeds were directly sown at the top of the rhizobox for *in-situ* germination. Plants were grown in rhizoboxes in a greenhouse at the King’s Building campus (Edinburgh, UK, 55°55’14.9”N, 3°10’09.9”W) and at the Debre Zeit Agricultural Research Centre, (Debre Zeit/Bishoftu, Ethiopia, 8°46’10.4”N, 38°59’55.6”E). Rhizoboxes were supported at 45° by metallic supports (Figure 1C), built using components described in Table S2. Rhizoboxes were placed in trays covered with anti-slip mesh. White polystyrene blocks were used as spacers between the support and rhizoboxes. The whole system was wrapped with sheeting to insulate against excessive radiative heating. Water was added to the trays at the base of the system and maintained to ensure a constant supply. The greenhouse day and night temperature setpoints were 26 and 16°C, respectively. These conditions were monitored throughout the experiment (Figure S6). See Supplemental Methods for more details.

### Imaging station

An imaging station for rhizoboxes was built (Figure 2) using components described in Table S3. The rhizobox is illuminated from the interior of the imaging station by two LED strips. An aluminium U-channel, parallel and medial to the rhizobox, supports the cameras. Five cameras were spaced 30 cm apart to ensure enough overlap between images for further stitching. The distance between lens and rhizobox was set at 78 cm. The imaging station was isolated from daylight using a black felt layer. See Supplemental Methods for more details.

### Cameras and image capture

We used the Phenotiki hardware platform (Minervini *et al.*, 2017), adapted for this project to allow for adjustable focus camera sensors (Raspberry Pi Camera) for imaging. Phenotiki sensor software was modified to trigger multiple cameras simultaneously by using a master-slave design. To reduce overhead during image acquisition, images were uploaded onto cloud-based storage (Google Drive) at scheduled times of the day (pipeline in Figure 3). (Alternatively data can be downloaded on request by the user from the master or uploaded to a local workstation should Internet access at the greenhouse be suboptimal.) Acquisition parameters (Supplemental Table S4) are the same for each device. To compensate for lens distortion, camera calibration was performed (Zhang 2000), using a chessboard of ArUco markers to determine the intrinsic camera parameters (Zhang 2000, Romero-Ramirez *et al.*, 2018). A series of permanent ArUco markers (4 cm^2^) were fixed to the interior of the imaging station frame to further improve picture assembly and also permit the co-registration of images acquired in longitudinal manner. This co-registration allows temporal analysis of growth at a local level (Figure 5, 6). See Supplemental Methods for more details.

### Image processing for stitching

Following image acquisition, image series of one rhizobox are processed to create a large mosaic stitching to obtain a single large image of the rhizobox (akin to the process of creating a panoramic image from multiple images). QR codes placed on the top corners of the glass pane on each rhizobox are decoded automatically from the stitched images to identify them. See Supplemental Methods for more details.

### Image segmentation

The stitched, large, image was used to segment the root system from soil background. Segmentation is performed in two steps: (*i*) Foreground-Background (FG-BG) segmentation; and (*ii*) noise removal. *FG-BG segmentation.* Imaging effects and artefacts preclude the use of a simple thresholding operation to separate root from soil. Therefore, we analysed the root images row by row to identify root pixels, which includes a parabolic threshold function to compensate for lateral illumination. *Noise removal.* Although the previous step is able to determine the plant roots, over-segmentation can still occur, due to clutter in the scene (e.g., presence of droplets inside the rhizobox). To alleviate this, we perform a refining step to remove the noise. Once the segmentation of the RSA is obtained, root traits are extracted as reported in Table 1. After the data were extracted from the segmentation mask, they were converted from pixels into cm. See Supplemental Methods and Figure S7 for more details.

### Local root density

To determine local root density the segmented image of a root system is sub-divided into a regular grid, where each cell is 200 x 200 pixels (ca. 9 cm^2^). For each cell, we compute the total number of root pixels from the segmentation mask and convert the measure to cm^2^. For dynamic analyses, we compute the difference of two consecutive root densities at time *t*_*n*_ and *t*_*n-1*_.

## Acknowledgements

The authors thank Joanna Jones and Şeyhmus Dündar for testing some initial rhizobox designs, Sam Anderson and Javier Surís Auguet for help with soil preparation and building in greenhouse, Peter Hoebe from Scotland’s Rural College (SRUC) for providing barley seeds. PD thanks BBSRC (BB IAA 15/16–PD & BB GC IAA 16/17) for funding which supported early-stage work with rhizoboxes. ST and PD thank BBSRC for funding (BB/P023487/1). CC thanks CONICYT PFCHA/DOCTORADO BECAS CHILE/2016 – 72170128 for a PhD scholarship. The authors declare no conflicts of interest.

## Author Contributions

All authors contributed to the development of the rhizobox growth system for visualising root growth, VG and ST developed and implemented image capture and data processing, TB, CC and IR developed the growth conditions for chickpea growth in rhizoboxes. TB, CC, VG, IR, ST and PD wrote the paper.

Thibaut Bontpart, 0000-0002-4243-8348

Cristobal Concha, 0000-0003-2724-2985

Valerio Guiffrida, 0000-0002-5232-677X

Ingrid Robertson, 0000-0001-6666-3876

Kassahun Admkie, 0000-0002-3535-1391

Tulu Degefu, 0000-0003-4962-8619

Nigusie Girma, 0000-0002-9322-9861

Kassahun Tesfaye, 0000-0002-4046-4657

Teklehaimanot Haileselassie, 0000-0001-5157-7086

Asnake Fikre, 0000-0003-3024-2070

Masresha Fetene, 0000-0003-0112-1210

Sotirios A. Tsaftaris, 0000-0002-8795-9294

Peter Doerner, 0000-0001-7218-8469

## Short supporting legends

**Table S1.** List of rhizobox components

**Table S2.** List of components for rhizobox support

**Table S3.** List of imaging station components

**Table S4.** Parameters for Phenotiki Sensor Setup

**Figure S1.** Correlation between chickpea root area and root length

**Figure S2.** Correlation between chickpea root area and solidity

**Figure S3.** Barley root system architecture parameters

**Figure S4.** Barley root area density and changes over time

**Figure S5.** Chickpea root systems in rhizoboxes tested in Ethiopia

**Figure S6.** Environmental conditions in greenhouse (Edinburgh, UK)

**Figure S7.** Segmentation method

**Movie S1.** Time-lapse development of root system in a rhizobox

