## Supporting Information for "Affordable and robust phenotyping framework to analyse root system architecture of soil-grown plants"

### Supporting Materials and Methods

#### Rhizobox design and construction

The rhizobox (Figure 1**A**, **B**), made of the components described in Supplemental Table S1, holds a 6 mm layer of soil between 150 cm high x 45 cm wide x 0.6 cm thick (external dimensions) glass pane as front and a backing sheet of polyvinylchloride (PVC) for a total soil volume of ~3.7 dm^3^. Two silicon strips are glued as spacers along the PVC sheet. The frame comprises two aluminium U-channels (inside lined with cloth tape) facing each other to close the rhizobox on the sides, and a wire inserted into a folded piece of nylon mesh to close the bottom of the rhizobox. The wire is inserted through a 2 mm hole positioned 5 mm from the base of each U-channel, and 5 cm extensions on each side are bent against the U-channel to hold it in place.

In Edinburgh, air dried and sieved (2mm aperture mesh) John Innes No.1 Young Plant Compost (Westland, UK) is homogenised with tap water to get 50% (w:w) water content, then sieved again (rotary sieve, 5mm aperture mesh) before loading into the rhizobox. After positioning the inner mesh at the base of the PVC sheet placed horizontally on a flat surface (silicon strips up), the soil is manually spread across then compressed using a hollow PVC roll (80 cm, 6 cm diameter), making sure that the soil is homogeneous and the surface is level with the silicon strips. After adding the glass pane, the system is closed with the frame described above.

In Ethiopia, around 95% of chickpea is grown and produced in vertisols. A field source for vertisol was identified at the EIAR in Debre Zeit (8°46'04.6"N 39°00'09.6"E) and around 100 kg of soil was collected from the site. This vertisol had 65-70% clay content. The soil was mixed regularly and sun-dried for several days until dry. The dry vertisol was sieved in a 1.5mm x 1.5mm size mesh sieve and around 60 kg was recovered. Subsequently the same protocol as in Edinburgh was followed.

#### Plant material and growth conditions

For UK experiments, chickpea (*Cicer arietinum* L., Kabuli type, Aldi, UK) seeds were imbibed by soaking in water with gentle shaking (roller mixer). After 2h, seeds were rinsed, left overnight in water, then sown in John Innes No.1 compost (50 % w:w) for 2-3 days before transplanting into a rhizobox. A hole was made in the soil at the top and centre of the rhizobox to insert a chickpea seedling root. This was immediately watered and covered in soil, then watered daily for five days to ensure plant survival. In Ethiopia two cultivars (*var.* ‘Dhera’, Kabuli; and *var.* ’Teketay’, Desi) were sown into small plastic pots filled with vertisol for germination and then transferred to the rhizoboxes 5d after germination. Barley (*Hordeum vulgare* L., variety *Concerto*, SRUC, UK) seeds were directly sown into the top of rhizoboxes for *in-situ* germination.

Plants were grown in rhizoboxes in a glass house at the King's Building campus (Edinburgh, UK, 55°55'14.9"N, 3°10'09.9"W) as described below and at the Debre Zeit Agricultural Research Centre, (Debre Zeit, Ethiopia, 8°46'10.4"N, 38°59'55.6"E). Environmental conditions in the Edinburgh glass house during the experiment shown in Figures 4-6 are shown in Supplemental Figure S6.

Metallic supports, designed to hold the rhizoboxes at 45° (Figure 1**C**), were built using components described in Table S2. Rhizoboxes were placed in trays covered with anti-slip mesh. White polystyrene blocks were used as spacers between the metallic support and rhizobox, and between each rhizobox. Once all rhizoboxes were in place, the whole system was wrapped with white-black sheeting (mylar sheet with white side facing outwards and black side facing inwards) to insulate against excessive heating. Water was added to the trays at the base of the system and maintained to ensure a constant supply. The greenhouse day and night temperature setpoints were 26 and 16°C, respectively. Temperature and air relative humidity were recorded using USB-502 data logger (Measurement Computing Corporation, USA; Figure S1).

#### Imaging station

A station for imaging rhizoboxes using a camera array was built in Edinburgh (Figure 2) using components described in Table S3. This structure was designed to keep the rhizobox at 15° angle during image capture to allow the rhizobox to be stable during imaging. The rhizobox is illuminated from inside the imaging station from each side by two LED strips turned on to the maximum intensity. They were glued inside an aluminium U-channel previously insulated with a white PVC strip and connected to a power supply with dimmer. Four pieces of aluminium trunkings were arranged in parallel inside the imaging station to hold the LED strip U-channels and allow fine tuning of their position. An aluminium U-channel, parallel and central to the rhizobox, was inserted into the imaging station as a support for the cameras. Five cameras were spaced 30 cm apart to ensure enough overlap between images for further stitching. The distance between lens and rhizobox was set at 78 cm. The imaging station was isolated from daylight by covering with black felt.

#### Camera and image capture

We used the affordable imaging hardware and software platform Phenotiki (Minervini *et al.,* 2017), after adapting it for this project to use adjustable focus camera sensors (Raspberry Pi Camera) for imaging. The Phenotiki sensor software was modified to trigger multiple cameras simultaneously by using a master-slave approach, where one Raspberry Pi (the master, configured with the extended Phenotiki Sensor Software) allows the other devices (the slaves) to connect *via* wireless communication. To acquire images, the master triggers and collects the images obtained from all the other devices and stores them locally. The user can operate the sensor *via* a web-based interface. To reduce overhead during image acquisition, images were uploaded into cloud-based storage (Google Drive) at scheduled times of the day (pipeline in Figure 3). However, in case of suboptimal connectivity, the user can also download the acquired images directly from the Phenotiki interface. Acquisition parameters (see Table S4) are the same for each device. Cameras are placed and configured in the imaging station. To compensate for lens distortion, camera calibration was performed (Zhang 2000), using a chessboard of ArUco markers (typically referred as ChAruco) to determine the intrinsic camera parameters (Zhang 2000, Romero-Ramirez *et al.,* 2018). A series of permanent ArUco markers (4 cm^2^) were fixed to the interior of the imaging station frame that flanks the aperture for the rhizobox to be visible by the cameras and further improve picture assembly. Images generated using one exposure were of sufficient quality across the horizontal extent of the rhizobox. Combining images from up to 5 different exposure settings in a “high dynamic range” (HDR) mode is also possible, but comes at the cost of increased computational load and storage requirements.

#### Image processing for stitching

Once an image series is acquired, it is processed for stitching to obtain a single large image of the rhizobox. First lens distortion and skew is removed (using the camera/lens parameters determined when the cameras were initially set up), then panorama stitching is applied (Brown and Lowe 2007). Specifically, this algorithm relies on the extraction of robust key points to compute pairwise image transformations (homographies). However, especially for the images where the root is not (yet) present (*e.g.*, early during the growth period where the roots are absent in images of the bottom of the rhizobox), distorted stitched images are further rectified using the permanent ArUco markers. The presence of the ArUco markers is also used to co-register the images, allowing a more robust temporal analysis of the RSA traits. QR codes placed on the top corners of the glass sheet on each rhizobox are decoded automatically from the stitched images to identify them and for accounting of experimental data. The final stitched image is cropped to only include the rhizobox in the field of view. After this operation, the final image size is approximately $9000\times2700$ pixels.

#### Image segmentation

Following image stitching, the large image was used to segment the root system from the soil background, which is an essential step to extract phenotyping data. Our RSA segmentation is performed in two steps: (*i*) Foreground-Background (FG-BG) segmentation; and (*ii*) noise removal.

*FG-BG segmentation.* Although roots have discernible contrast with respect to soil, clutter and artefacts such as LEDs illuminating a few water droplets in the soil, resulting in high intensity pixels, preclude the use of a simple thresholding operation to separate root from soil. Therefore, we devised a different approach that analyses the root images row by row to identify root pixels (Supplemental Figure S7). Specifically, the stitched image $I$ of size $W\times H$ is converted into grayscale and smoothed with a Gaussian filter to reduce clutter. Then, the root image is scanned from bottom up, where each row$u$ in $I_{u}(v)$ is analysed at each iteration, as shown in Supplemental Figure S7**A**. In our algorithm, $I_{u}\left( v \right)$ represents the intensities of the image at row $u$, as plotted in Supplemental Figure S7**B**. In order to isolate roots from soil, we model the background with a function $\varphi_{u}\left( v \right)$ (details how this function is determined are provided below). Intuitively, we remove for each line the background $\varphi$ from $I$ for each row $u$, and all the pixels above a threshold are considered as root. Formally, this process to obtain segmented image $\hat{I}_{u}(v)$ is defined as follows:

$\hat{I}_{u}\left( v \right)=\left( I_{u}\left( v \right)-\varphi_{u}(v) \right)\geq\tau\left( v \right)$,

where $\tau\left( v \right)$ is a threshold function that varies with respect to the horizontal coordinate $v$ (Supplemental Figure S7**C**). The function $\varphi_{u}(v)$ is computed as the average intensity of all the pixels below $u$, excluding all root pixels segmented up to that point. Formally, the background function is defined as:

$$\varphi_{u}\left( v \right)= \frac{1}{H-u}\sum_{u\in I\setminus\hat{I}} I_{u}(v),$$

For the function $\tau\left( v \right)$, we selected a parabolic threshold function to compensate for the lateral illumination with LEDs that results in pixels towards the sides of the rhizobox to be brighter. For our data, we computed $\tau(v)$ such that $\tau\left( \frac{W}{2} \right)=65$ (the parabola vertex) and $\tau\left( 0 \right)=\tau\left( W \right)=85$. This approach is inspired by the parametric approach used by Tsaftaris and colleagues that corrected for a non-uniform distortion in the presence of objects of interest in the context of Atomic Force Microscopy (Tsaftaris *et al.,* 2008a, Tsaftaris *et al.,* 2008b).

*Noise removal.* Although the previous step is able to determine the plant roots, over-segmentation can still occur, due to clutter in the scene (*e.g.*, presence of droplets inside the rhizobox). In order to alleviate this, we perform a refining step to remove the noise. Firstly, small isolated white pixel regions are removed from the segmentation mask, when either the number of pixels is equal or less than 15, or it exhibits a round shape (stockiness^[1]^ > 0.75). Measuring the roundness is important, as small root regions are characterized by low stockiness instead. However, this procedure does not remove all the cluttered areas in the segmented image and a further post-processing step is necessary. Generally speaking, RSAs should be the largest connected component in a segmented image. In our case, due to gaps caused by soil occlusions, RSA is composed by several connected components, which are hard to isolate from clutter. Therefore, we identify which is the densest area. Specifically, we compute the number of segmented pixels in non-overlapping regions of $70\times70$ pixels, overcoming gaps in the RSA. At this point, we select as root the densest area and other (cluttered) areas are discarded.

Once the segmentation of the RSA is obtained, root traits are extracted as reported in Table 1. After the data are extracted from the segmentation mask, they were converted from pixels into cm.

#### Local root density

To determine local root density the segmented image of a root system is sub-divided into a regular grid, where each cell is $200\times200$ pixels. This corresponds to an area of approximately 9 cm^2^. For each cell, we compute the total number of root pixels from the segmentation mask and convert the measure to cm^2^. From the root densities, we can also determine which parts of the root system are growing more with respect to others by computing the difference of two consecutive root densities at time $t_{n}$ and $t_{n-1}$.

### Supporting data

- **Table S1**. Rhizobox components
- **Table S2**. Support components
- **Table S3**. Imaging station components
- **Table S4**. Phenotiki Sensor Setup
- **Figure S1**. Correlation between root area and root length
- **Figure S2**. Correlation between root area and solidity
- **Figure S3**. Barley root system architecture parameters.
- **Figure S4**. Barley root area density.
- **Figure S5**. Chickpea root systems in rhizoboxes tested in Ethiopia.
- **Figure S6**. Environmental conditions in glass house (Edinburgh, UK).
- **Figure S7**. Segmentation method.
- **Movie S1**. Time-lapse development of root system in a rhizobox.

**Table S1**. Rhizobox components.

| Component | Dimensions (cm): Length x Width x Thickness | Number | Price (£)* | Supplier (City, Country) |
| --- | --- | --- | --- | --- |
| Glass sheet | 150 x 45 x 0.6 | 1 | 51 | Alba Glass (Musselburgh, UK) |
| PVC sheet | 150 x 45 x 0.6 | 1 | 27 | Direct Plastics (Sheffield, UK) |
| Silicon strips | 150 x 1 x 0.6 | 2 | 17.71 | Silex silicones Ltd (Bordon, UK) |
| Inner nylon mesh | 45 x 5,  8 layers of 0.1 aperture mesh | 1 | 0.74 | Buzzstop (Romiley, UK) |
| Outer nylon mesh | 45 x 3,  8 layers of 0.1 aperture mesh | 1 | 0.46 |  |
| Aluminium U-channel | External: 150 x 2.5 x 0.3  Internal: 150 x 1.9 x 0.3 | 2 | 13.9 | Aluminium warehouse (Hatfield, UK) |
| Cloth tape | 150 x 1.9 x 0.033 | 4 | 0.96 | Advance Tapes (Thurmaston, UK) |
| Wire | 55, diameter: 0.16 | 1 | 0.67 | JAC tools (Aylesbury, UK) |
| Total |  |  | 112.44 |  |

* Calculated dividing the price of the batch (including cutting, VAT and delivery) by the number/quantity necessary to build one rhizobox.

**Table S2**. Support components.

| Component | Dimensions (cm):  Length x Width x Thickness | Number | Price (£)* | Supplier (City, Country) |
| --- | --- | --- | --- | --- |
| Dexion slotted angle DX160 | 304.8 x 4 | 8 | 112.18 | Racking man (Leeds, UK) |
| Punched Strap | 152.4 x 3.8 | 8 | 13.7 |  |
| Metal Footplates |  | 2 | 3.35 |  |
| Bolts and nuts |  | Pack of 100 | 7.9 |  |
| Giant Garden Trays | 110 x 55 x 4 | 2 | 21.16 | Garland (Kingswinford, UK) |
| Anti-slip mesh | 150 x 50 | 2 | 13.98 | Amazon |
| Polystyrene sheets, ECO100 grade | 120 x 60 x 6 | 20 | 183.38 | Custompac (Castleford, UK) |
| Total |  |  | 355.65 |  |

* Calculated dividing the price of the batch (including cutting, VAT and delivery) by the number/quantity necessary to build one support holding 18 rhizoboxes (2 rows of 9).

**Table S3**. Imaging station components

| Component | Dimensions (cm) | Number | Price (£)* | Supplier (City, Country) |
| --- | --- | --- | --- | --- |
| Dexion slotted angle DX160 | 304.8 | 12 | 184.52 | Racking man (Leeds, UK) |
| Metal Footplates |  | 8 | 11.50 |  |
| Bolts and nuts |  | Pack of 100 | 7.9 |  |
| Aluminium trunking | 92 | 4 | Already available | Polybuild (Washington, UK) |
| Aluminium U-channel | External: 150 x 2.5 x 1.9cm;  Internal: 150 x 1.9 x 1.6  Thickness: 0.3 | 3 | 20.92 | Aluminium warehouse (Hatfield, UK) |
| Aluminium Angle | 5 x 5 x 0.3 | 6 | 19.2 |  |
| Feet |  |  | Already available |  |
| LED strip (natural white colour, 4.8 W.m^-1^) | 150 | 2 | 17.41 |  |
| LED power supply with dimmer (12 V, 2 A) |  | 1 | 25.09 | Powerpax (UK) |
| UK plug |  | 1 | 1.8 | Pro-elec (UK) |
| Black felt | 10 m x 150 cm | n/a | 44.85 | Fabric land (UK) |
| Raspberry PI 3 Model B V1.2 |  | 5 | 176.35 | Farnell (UK) |
| 16GB MicroSD Card with preloaded OS Installer |  | 5 | 65.7 |  |
| Official Raspberry Pi International PSU |  | 5 | 41.35 |  |
| Case for Raspberry Pi 3 |  | 5 | 41.35 |  |
| Cameras (Raspberry Pi 3 model B V1.2) |  | 5 | 66.9 | Waveshare (US) |
| Total |  |  | 724.84 |  |

* Calculated dividing the price of the batch (including cutting, VAT and delivery) by the number/quantity necessary for the imaging station.

**Table S4.** Acquisition parameters as configured for all the Phenotiki sensors to acquire root images.

| Parameter | Value |
| --- | --- |
| Resolution | 3280x2464 |
| ISO | 800 |
| Exposure Mode | Night Preview |
| Automatic White Balance (AWB) | Tungsten |

#### Supplemental Figure Legends

**Supplemental Figure S1**

**Correlation between RSA parameters.** The correlation between root length and root area of chickpea plants in rhizoboxes was plotted.

**Supplemental Figure S2**

**Correlation between RSA parameters.** The correlation between root length and solidity of the chickpea root system of plants grown in rhizoboxes was plotted.

**Supplemental Figure S3**

**Barley RSA parameters.** Evolution of root system architecture parameters of a barley (*Hordeum vulgare*, *var.* Concerto) in a rhizobox. All parameters (root system depth, width, convex hull area, centroid, relative growth rate, length, area and solidity) were extracted from images collected at 8, 11, 15, 18, 22 and 25 days after sowing.

**Supplemental Figure S4**

**Root area density analysis of barley.** Computed from root pixels in a grid of 200 x 200 pixel squares covering the whole rhizobox. The days after sowing (das) are indicated on panels. The scale unit corresponds to the number of root pixels per square. (**A**) Absolute root area density. (**B**) The rate of root area density change was computed as the difference of root pixels between time *t_n_* and time *t_n-1_*, over the difference in time ($\Delta$das).

**Supplemental Figure S5**

**Figure S5**. Chickpea root systems in rhizoboxes tested in Ethiopia. Top row (A1-C1): three different plants growing in rhizoboxes filled with vertisol. Bottom row (A2-C2): Segmented images corresponding to the root systems shown in A1-C1, respectively.

**Supplemental Figure S6**

**Environmental conditions in Edinburgh glass house.** Data were collected during the growth in rhizoboxes of the four chickpea plants from which root system architecture parameters were extracted. Air temperature and relative humidity were automatically recorded every hour with a USB-502 data logger (Measurement Computing Corporation, Norton, USA) suspended at the level of the chickpea canopy. The shaded part corresponds to day time according to local sunrise and sunset. Violin plots (<http://shiny.chemgrid.org/boxplotr/> using the vioplot package <https://cran.r-project.org/web/packages/vioplot/index.html> ) show computed median (open circle) extent of first and third quartiles (thick lines) and whiskers (thin lines).

**Supplemental Figure S7**

**Root segmentation algorithm.** (A) The image of a root is scanned line by line from bottom to the top. Each line $u$ is represented as $I_{u}(v)$, whereas $\varphi_{u}\left( v \right)$ is the function representing the background information, computed as the average of all the pixels below $I_{u}\left( v \right)$ excluding root pixels that might have been already segmented. (B) Plot of the line $I_{u}\left( v \right)$ (blue) and the background function $\varphi_{u}\left( v \right)$ (in orange). (C) Difference between the current line and the background (in blue) and the parabolic threshold function (in orange).

**Movie S1**. Time-lapse development of root system in a rhizobox.

1. Roundness (or stockiness) is computed as $4\pi\frac{\text{Area}}{\text{Perimeter}}$
