## Supplemental Figures for "Affordable and robust phenotyping framework to analyse root system architecture of soil-grown plants"

Supplemental Figure S1

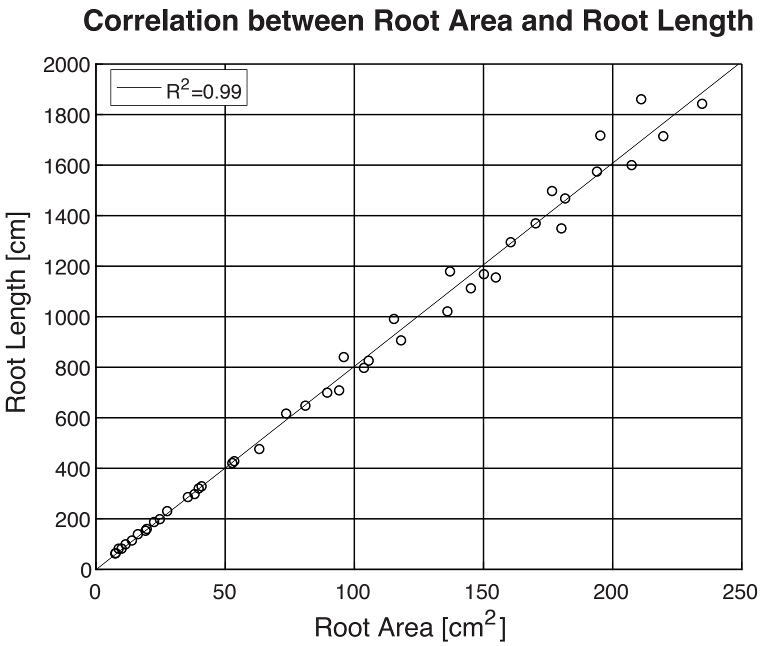

Supplemental Figure S2

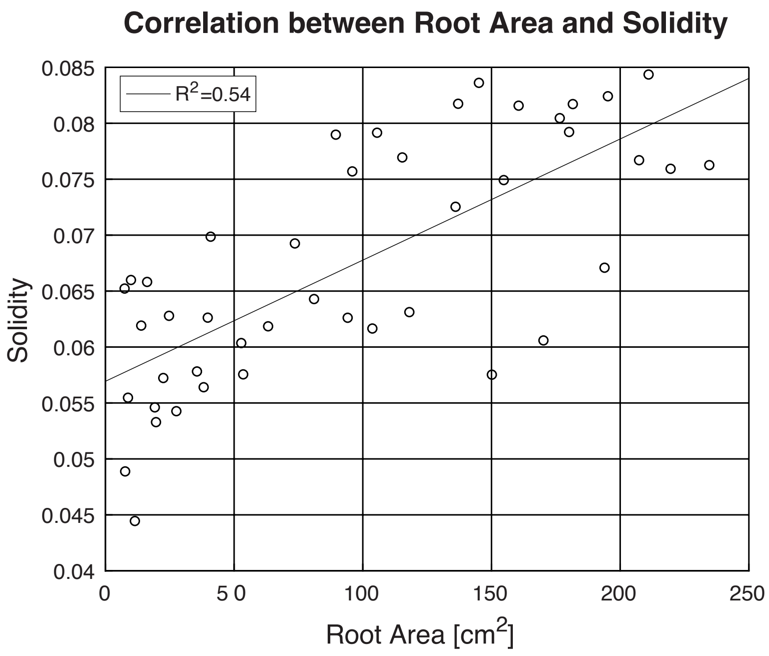

Supplemental Figure S3

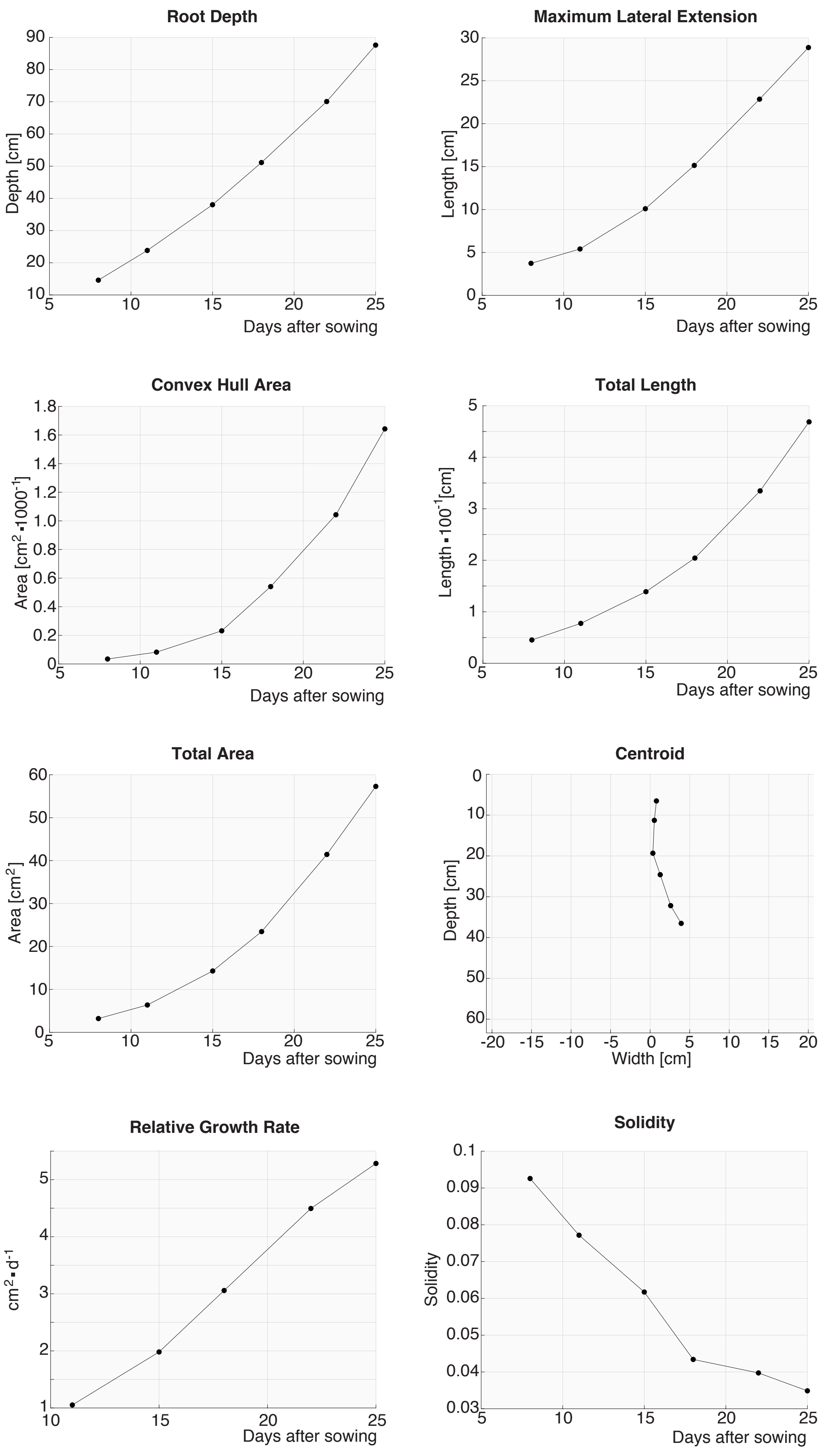

Supplemental Figure S4

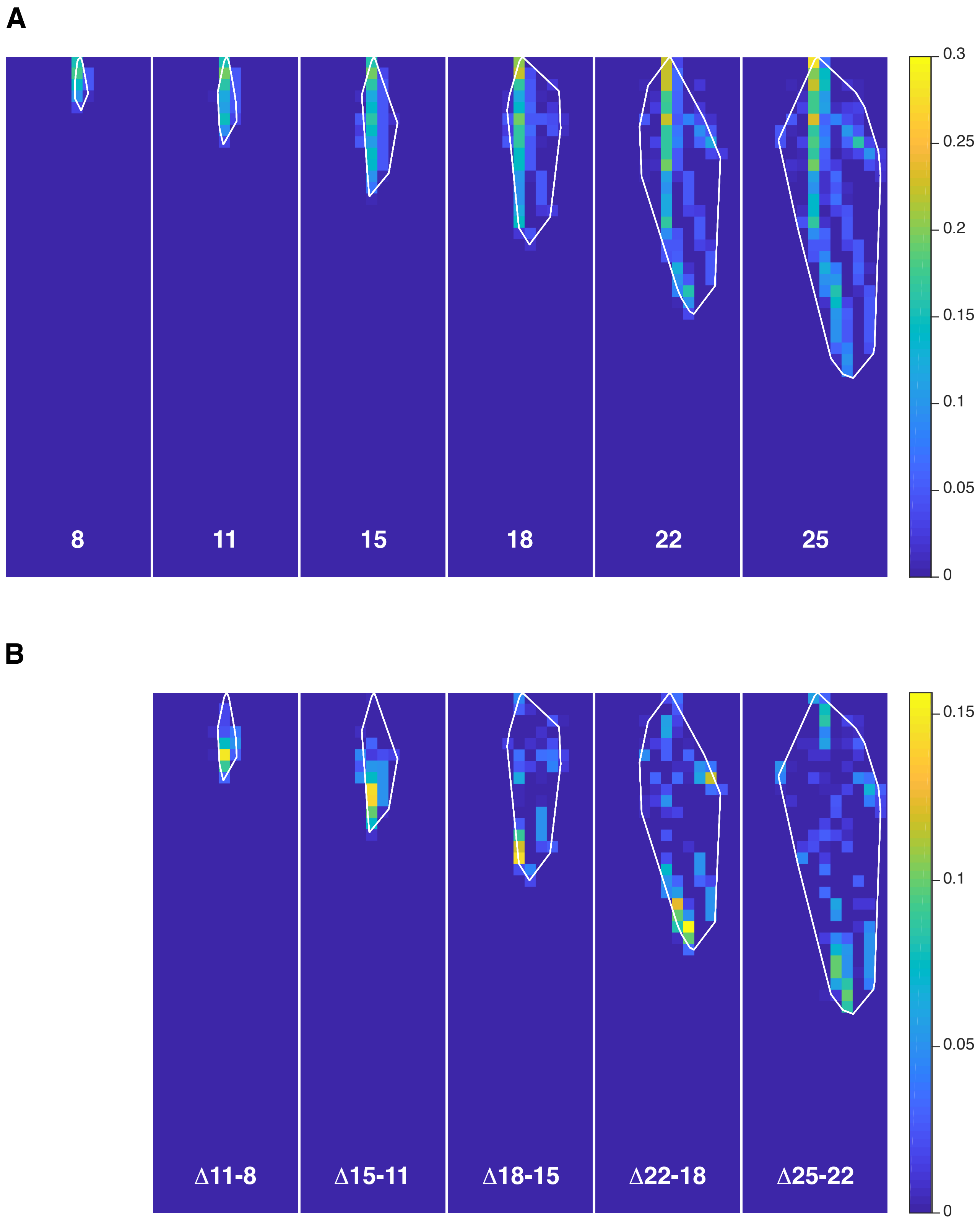

Supplemental Figure S5

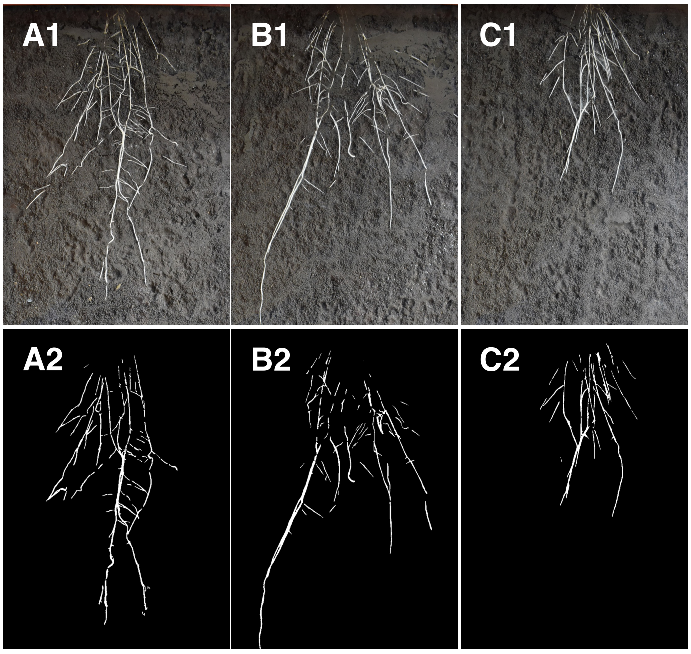

Supplemental Figure S6

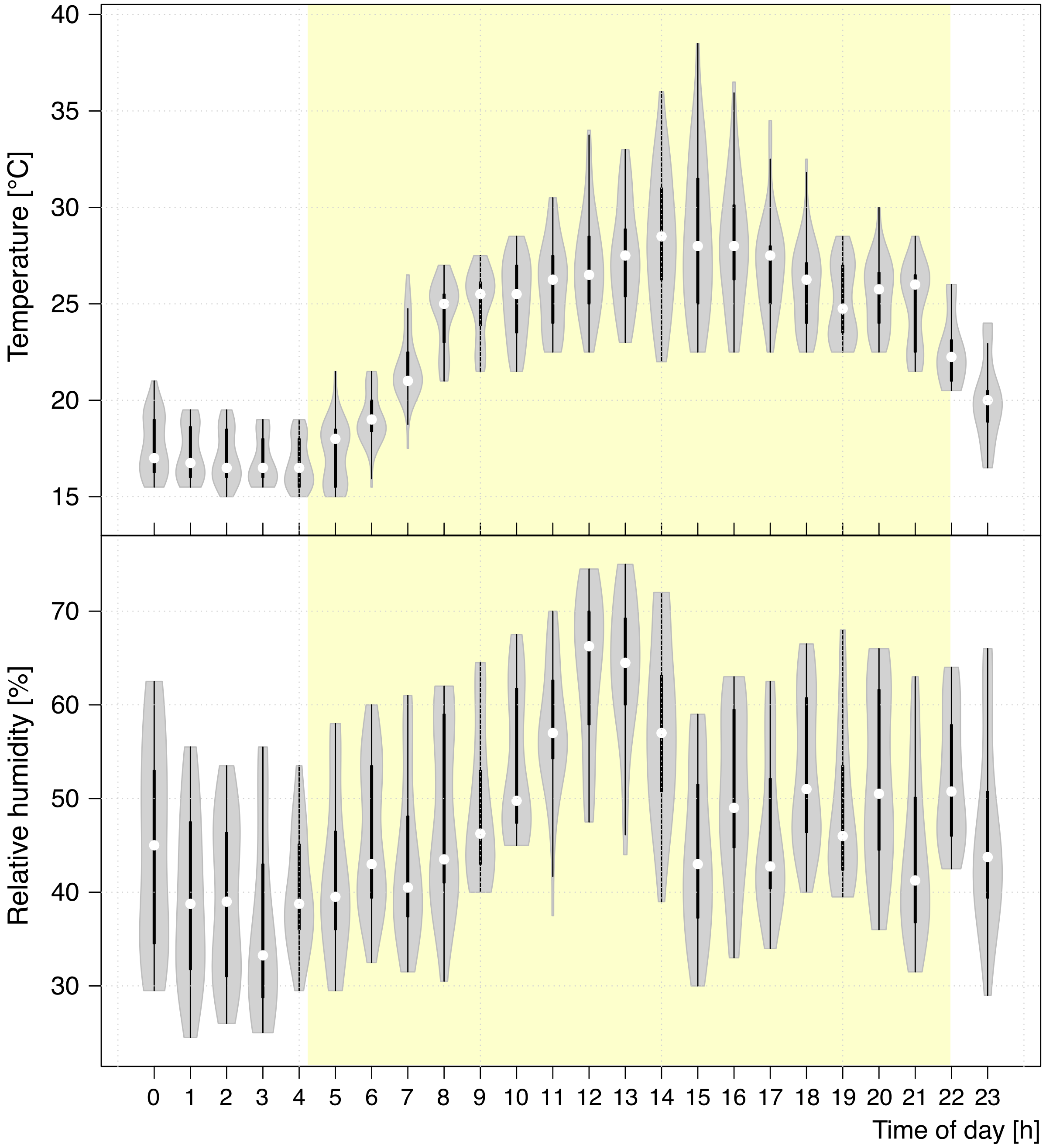

Supplemental Figure S7

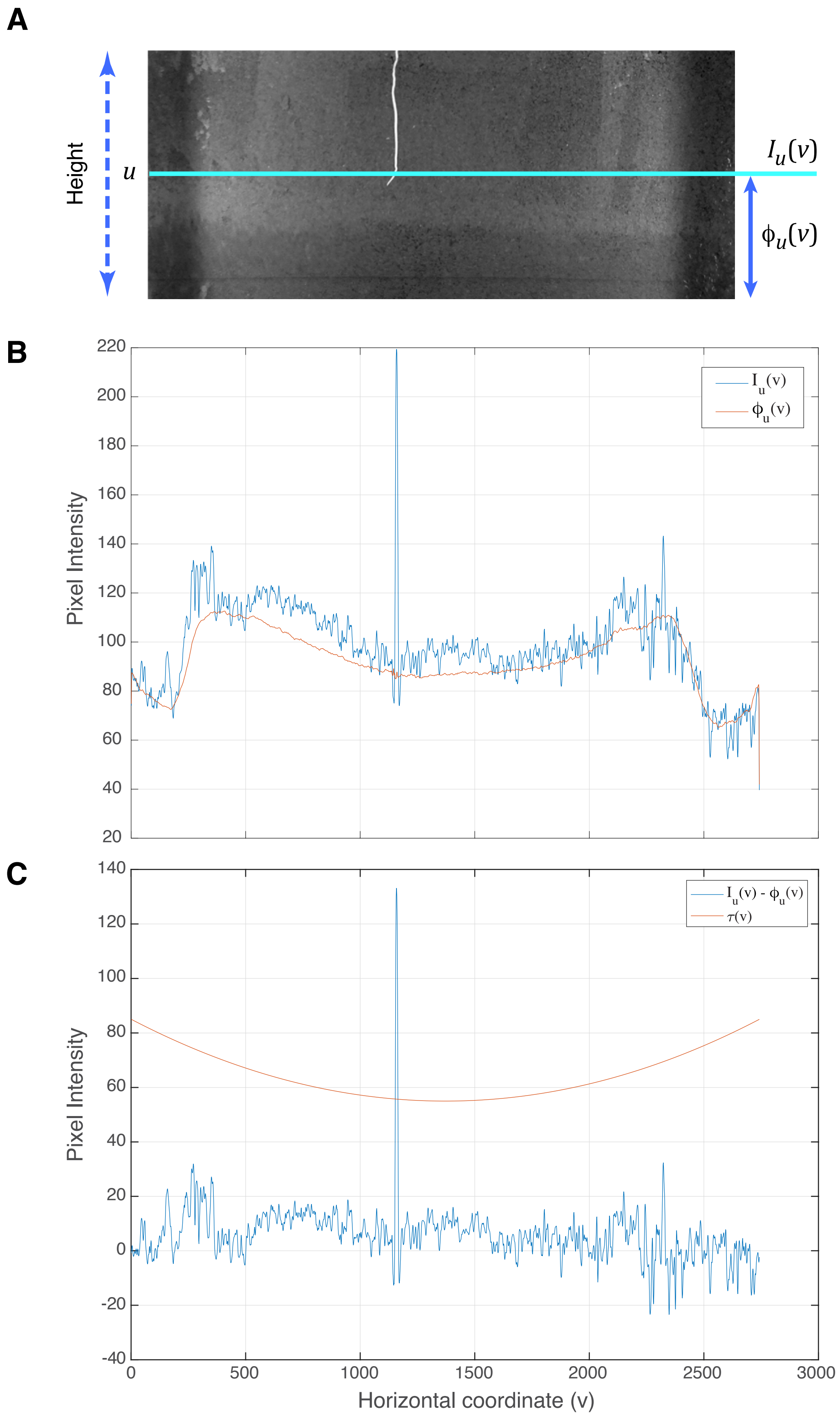
